# Mutation in DsbA signal sequence hampers the SRP mechanism: A new strategy to combat virulence factor

**DOI:** 10.1101/261537

**Authors:** Faiza Gul Durrani, Roquyya Gul, Muhammad Usman Mirza, Naheed Nazly Kaderbhai, Mahjabeen Saleem, Muhammad Waheed Akhter

## Abstract

Disulphide bond (Dsb) protein, characterized as an important virulence factor in gram negative bacteria. In this study, amino acid mutations in DsbA signal sequence (ss) and its effect on translocation of recombinant Ovine growth hormone (rOGH) was observed. Eight constructs were designed on the basis of increased hydrophobicity and showed that hydrophobicity and specificity of amino acid plays a crucial role in translocation of rOGH. Two DsbAss with the same hydropathy (1.539), one had alteration at -13 and second at -11 position; alanine (Ala) to isoleucine respectively were designed. The former DsbAss translocated rOGH from membrane to cytoplasmic fraction in *E. coli* as confirmed by SDS-PAGE, Western blot and molecular modelling analysis. MD simulations and binding free energy calculations evidenced that, altering Ala changed the orientation of signal peptide in the Ffh-M domain binding groove and hampered the process of translocation while change at position -11 pointed it outward. We hypothesize, amino acid and position of mutations in DsbAss can hinder the translocation process of signal recognition particle system, thus affecting the virulence of bacteria.

## Introduction

One of the main focus of scientists nowadays in drug design is to find the possible ways to disarm virulence factors of bacteria. This approach is needed as we enter the “post-antibiotic era” where all previously common treatable infections cannot be treated with antibiotics. The scientists predict that if they do not find new ways to kill the drug resistant pathogens by 2050, the annual mortality rate linked with multi drug resistant pathogens will surpass that of cancer, with 10 million deaths annually [1]. Therefore, new strategies are being developed to inhibit bacterial virulence rather than bacterial growth [2, 3]. Most of these virulence factors needs disulphide bonds to make them functional [4]. That is why the new research is focused on central system that regulates the deployment of multiple virulence factors in bacteria; the disulphide bond (Dsb) oxidative folding machinery [5, 6]. The most taxonomically widespread Dsb protein is DsbA, which is found in all classes of Proteobacteria and Chlamydiales along with numerous species of Fusobacteria and `Actinobacteria [4, 7]. Bacterial cells containing DsbA null mutations show a pleiotropic phenotype due to the incorrect folding of many periplasmic proteins (alkaline phosphatase, β-lactamase and Outer membrane protein A) and reduced fitness in animal models [8, 9]. Furthermore, they display attenuated virulence since the folding, stability and function of many bacterial virulence factors including toxins, secretion systems, adhesions and motility machines are dependent upon DsbA/DsbB mediated disulphide bond formation [4, 7]. For example, *V. Cholerae*, Enteropathogenic *E. coli* and Bordetella pertussis DsbA mutants secrete reduced levels of cholera, heat-labile and pertussis toxins respectively [10, 11].

We hypothesize a new strategy to hamper the Dsb system, which is to study the mutations in DsbA signal sequence (ss) rather than finding inhibitors for DsbA and observe its effects on translocation of recombinant ovine growth hormone (rOGH). The rOGH is a 22kDa protein and consists of 2 disulphide bonds. We have already reported the successful yield and translocation of soluble rOGH into the inner membrane of *E. coli* by using DsbAss and T7 promoter system [12]. The DsbAss works with signal recognition particle (SRP) targeting mechanism and has been the best choice for translocation of recombinant proteins to the inner membrane of *E. coli* [13]. The SRP mechanism in *E. coli* constitutes a “fifty-four homologue” (Ffh, also called P48) protein, necessary for viability and efficient protein export, and 4.5S RNA, respectively. The primary sequence of Ffh is divided into three domains: the N-domain residing at the amino terminus; G-domain, which harbours the GTPase activity and a methionine-rich, M-domain involved with SRP54 binding of both RNA and the signal sequence [14]. The interaction of Ffh with the signal sequence increases with the hydrophobicity of the sequence. It has been suggested that the magnitude of the interaction between SRP and the signal sequence correlates with the translocation efficiency as precursors with a more hydrophobic signal sequence are transported more efficiently [15, 16]. The Ffh’s structural insight gives an adequate understanding of the signal sequence recognition mechanism via SRP [15, 17]. The M domain of SRP54, through a series of functional and cross-linking studies [18-22] has been identified to comprise the signal sequence binding site. Successive events during the SRP cycle require a rearrangement of the relative position of the N, G and M domains. The binding of signal sequence within M domain induces conformational changes which stimulates the rotation of N, G domain to bring the GTPase domain closer and proceed with the translocation process [23, 24]. The structures provide evidence for a coupled binding and folding mechanism in which signal sequence binding induces the concerted folding of the GM linker helix, the finger loop, and the C-terminal alpha helix. This mechanism allows for a high degree of structural adaptability of the binding site and suggests how signal sequence binding in the M domain is coupled to repositioning of the N, G domain [17, 23-26].

In this study we explored the effect of amino acid substitution in all three parts of DsbAss; specifically hydrophobic (H) domain and its role in SRP targeting mechanism. In this investigation, the N, C and specifically H region of DsbAss were mutated and the effect on the translocation of the rOGH in *E. coli* was observed. The study also aimed to understand the structural insight of signal peptide conformation inside the Ffh-binding groove by using molecular dynamics (MD) simulations, molecular mechanics generalized born surface area (MM-GBSA) binding free energy calculations and to find the decisive residues with good interaction energies through per-residue decomposition analysis of important amino acids lining the hydrophobic binding groove. Here we report the dynamics behaviour of change of amino acids in the H-domain of signal peptide, in good agreement with the experimental data.

## Results

### rOGH expression analysis

The *E. coli* BL21 (DE3) codon plus cells transformed with pOGH-1 to -8 variants were cultured in LB-ampicillin medium at 37°C to an OD_600_ of 0.5, followed by induction with 10µM IPTG for 6 hrs. Equal amount of cells (based on OD_600_ values) were lysed from each culture and the protein expression was analysed by 15% SDS-PAGE. The expected molecular mass of rOGH is ∼22kDa, whereas variations were observed both in mass and expression levels of the total protein lysates of the *E. coli* transformed with different pOGH constructs **(Fig 1)**. As shown, the pOGH constructs -1, -2, -6 and -7 showed band of rOGH at the expected position i.e., ∼22kDa on the SDS-gel whereas the pOGH 3, -5 and -8 showed the band of expressed proteins at positions a little higher (∼25kDa) than the calculated molecular mass of rOGH. Since, the approximate size of 18 amino acids long DsbAss is 2kDa, the higher molecular mass species of rOGH in these constructs was likely to be the result of incomplete processing of DsbAss.

**Fig 1:**
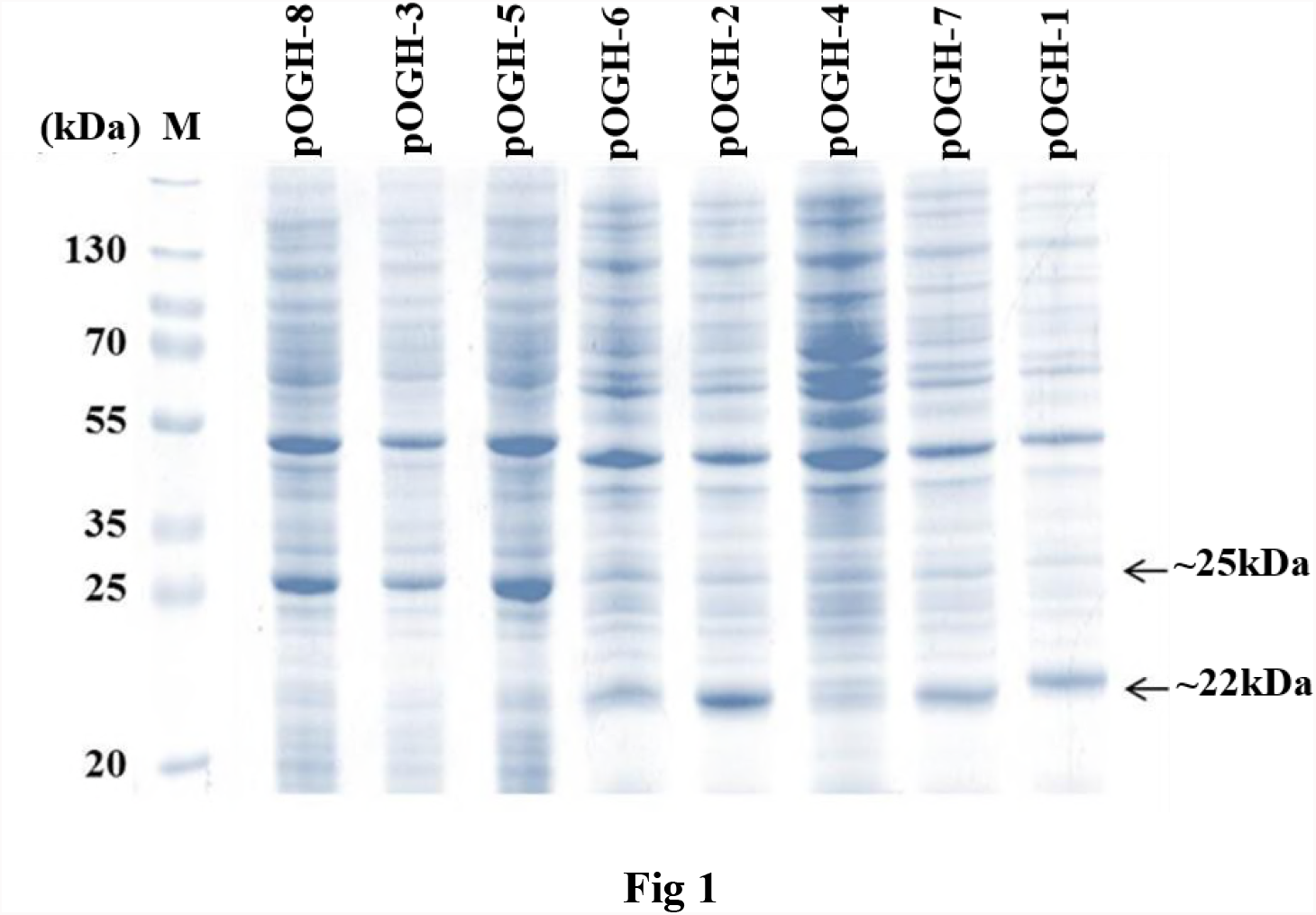
Total cell protein analysis of *E. coli* transformed with pOGH constructs in 15% SDS-gels. M represents molecular weight marker. pOGH -1 to -8 showing expression of rOGH in eight constructs.

In the case of pOGH-4 construct, size of the expressed rOGH was 22kDa but the expression level was barely detectable on SDS-gel and was confirmed by western blot analysis (data not shown).

### Impact of Ala substitutions in DsbAss H-domain

The DsbAss has four Ala residues in its H and near H-domain region and are present at position -3, -6, -11 and -13 with respect to the signal peptidase cleavage site **(Table 2)**. In the present study, these Ala residues were replaced by Ile in pOGH -2 to -6 constructs and impact of the substitutions was observed on rOGH expression and export in *E. coli*. Expression levels of rOGH in these constructs analysed by SDS-PAGE, ranged from undetectable (rOGH-4) to up to 25% (rOGH-5) of the total *E. coli* cellular proteins. Variations in the molecular mass were also observed in the case of rOGH-3 and -5 (∼25kDa) and rOGH-2, -4 and -6 proteins (∼22kDa).

**Table 1:**
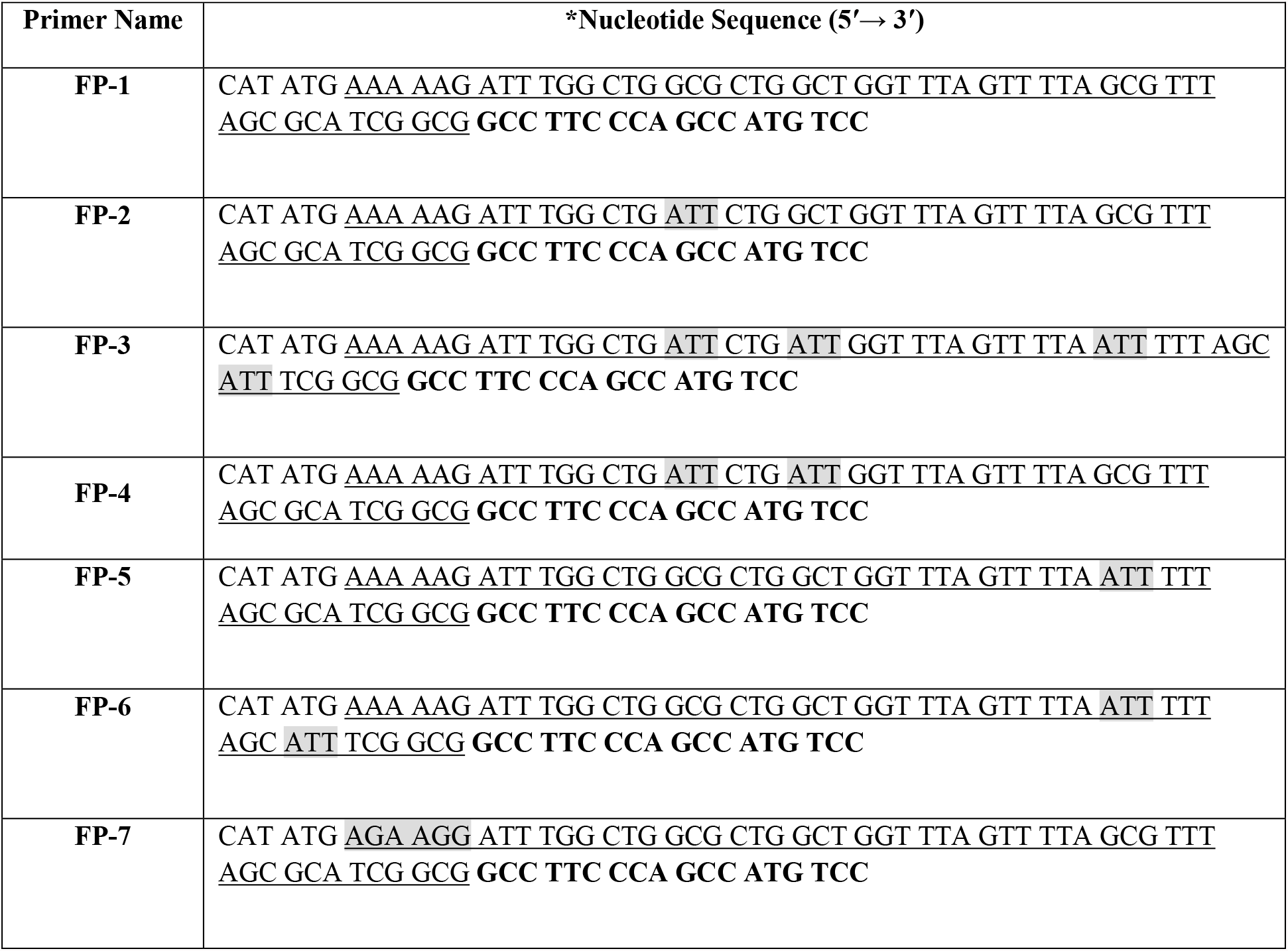
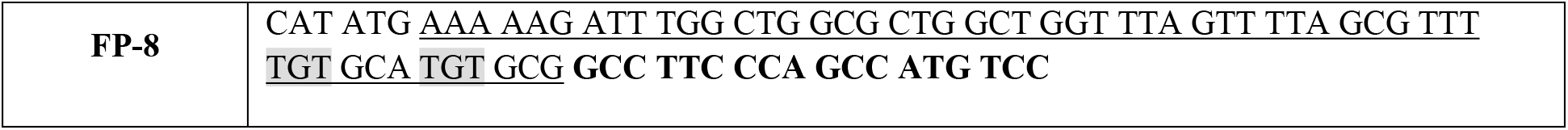
Sequence of primers used for construction of pOGH -1 to -8 plasmids. *The underlined sequence is DsbAss whereas bold represents the OGH cDNA sequence and non-underlined sequence is the site for *Nde* I restriction enzyme. Nucleotide variation incorporated in native DsbAss are highlighted in grey.

**Table 2:**
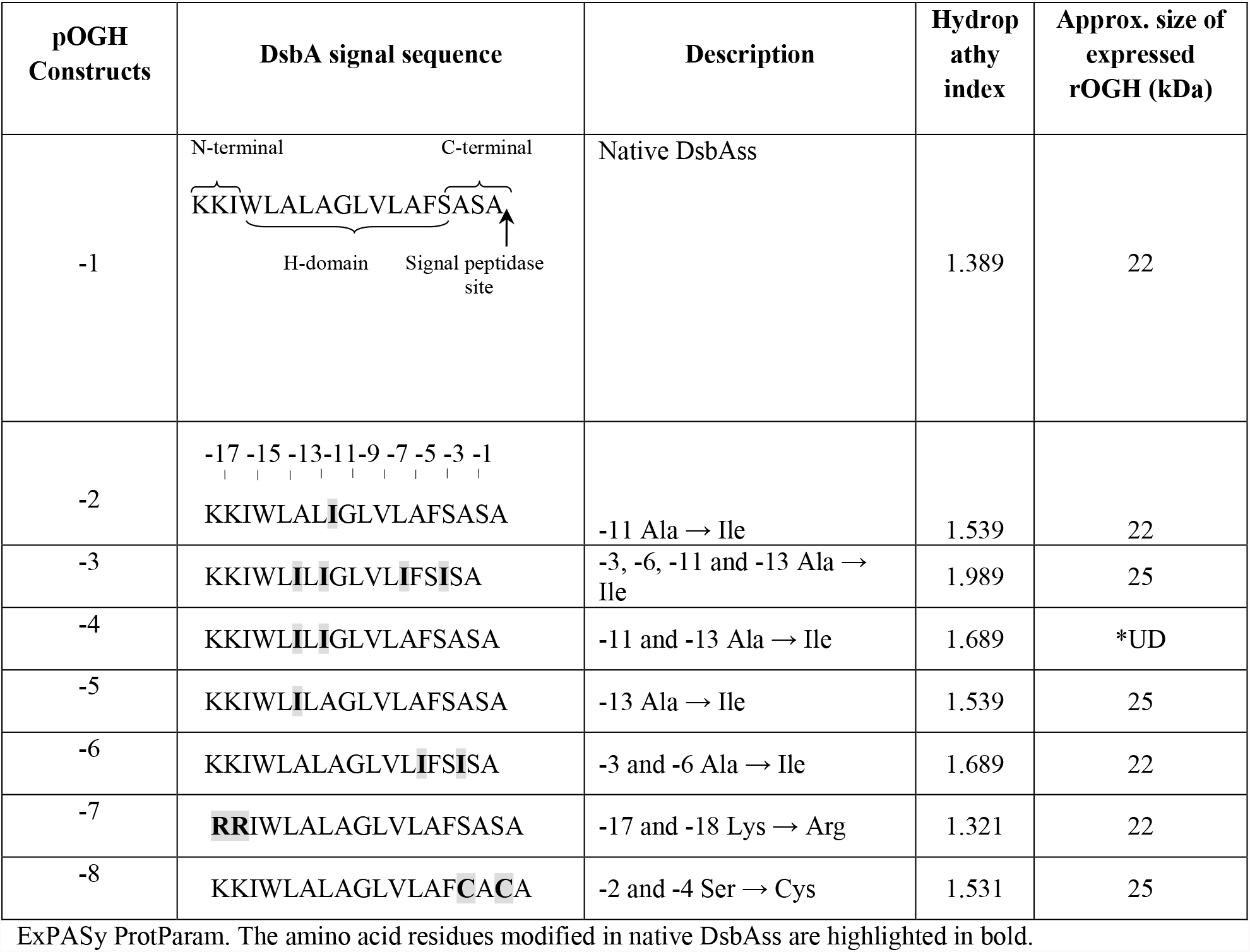
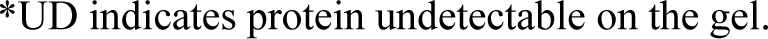
Hydropathy indices of native and modified DsbAss in pOGH constructs calculated using Swiss

In order to understand the behaviour of expressed rOGHs, the hydropathy index of each DsbAss mutant construct was analysed using the Swiss ExPASy Protparam tool **(Table 2)**. The pOGH -2 and -5 proteins had the same hydropathy indices but variable molecular weight of expressed rOGH and were of most interesting constructs. The sub cellular fractionation of proteins from the cells transformed with these constructs showed that in case of pOGH -2, DsbAss directed rOGH into the inner-membrane of *E. coli* **(Fig 2A)**. While in the case of pOGH -5, more than 75% of the expressed protein remained in the cytoplasmic fraction with negligible traces in the membrane fraction **(Fig 2B)**. Thus, Ala13 of DsbAss when substituted with Ile13 somehow hampered the export and processing of rOGH in *E. coli*. This suggests that it is not the hydropathy but the nature of amino acid substitution at specific position, which influences the mechanism of protein translocation. On the basis of this finding a model was hypothesized showing that by altering Ala at position -13 in hydrophobic region of DsbAss there is some conformational change which hampers the process of translocation. Therefore the protein which is destined for membrane fraction does not reach the inner membrane **(Fig 3)**.

**Fig 2:** **(A)** Analysis of sub-cellular protein fractionations of *E. coli* harbouring pOGH -2 by 15% SDS-PAGE and Western blotting. Lane 1: Un-induced sample; Lane 2: Total cell protein fraction from induced cells; Lane 3: Cytoplasmic fraction; Lane 4: Periplasmic fraction; Lane 5: Membrane fraction; Lane 6: Soluble fraction; Lane 7: Western blot of the purified rOGH -2 (Arrow indicates the position of rOGH at ∼22kDa). **(B)** Analysis of sub-cellular protein fractionations of *E. coli* harbouring pOGH -5 by 15% SDS-PAGE. Lane 1: Membrane fraction; Lane 2: Standard rOGH showing band at ∼22kDa; Lane 3: Periplasmic fraction; Lane 4: Cytoplasmic fraction; Lane 5: Total cell protein fraction from induced cells; Arrow indicates the position of rOGH at ∼25kDa.

**Fig 3:**
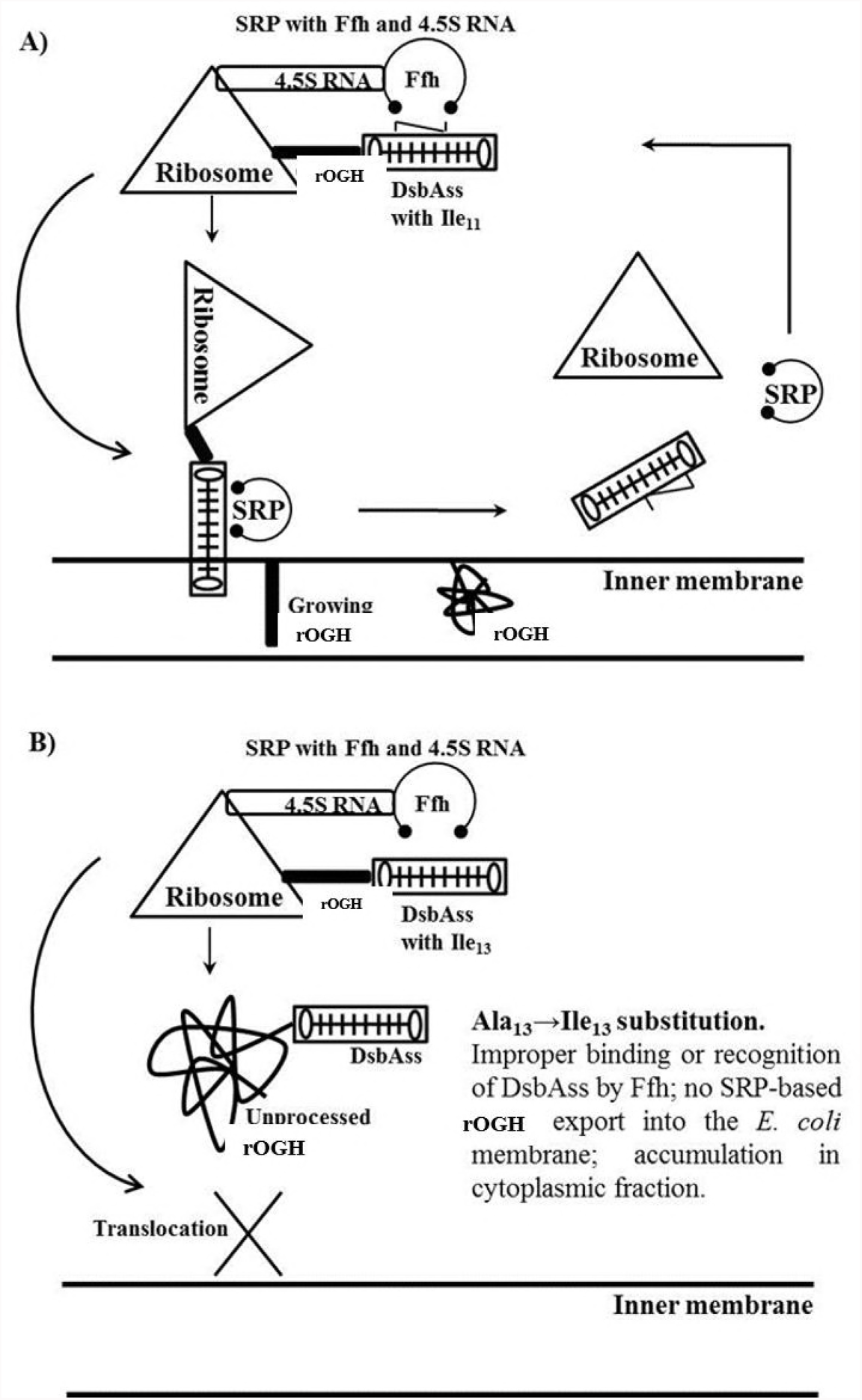
A model for SRP routing of rOGH through DsbAss. **(A)** DsbAss with Ala_11_→Ile_11_ substitution; **(B)** DsbAss with Ala_13_→Ile_13_ substitution.

### pOGH construct with Lys → Arg substitution at the N terminal of DsbAss

The two Lys residues of the N-terminal domain of DsbAss were substituted with Arg (pOGH -7 construct), as the presence of Arg in the N-region has previously been reported to be associated with better protein export [27]. Expression and translocation of recombinantly expressed protein was analysed by SDS-PAGE **(Fig 1)** and shows that the molecular mass of rOGH-7 was ∼22kDa and the protein was found in the membrane fraction indicating the complete process of the DsbAss. However, the presence of Arg in the N-terminal domain did not seem to result in enhanced rOGH expression and/or its export into the *E. coli* inner membrane.

### Influence of Ser → Cys substitution (at C terminal of DsbAss) on rOGH translocation

In pOGH -8 construct, two Ser residues present in the C-terminal domain of DsbAss were substituted with two Cys residues. Presence of Cys in the DsbAss has previously been described to be associated with the formation of disulphide bonds in the *E. coli* periplasm due to the oxidation-reduction role of DsbA [28]. In the present study Ser → Cys substitution affected the translocation process of rOGH as analysed by SDS-PAGE and was found in the cytoplasmic fraction with a molecular mass of ∼25kDa **(Fig 1)**. Apparently, this substitution changed the polar properties of C-terminal region, which affected the cleavage of DsbAss and led to the accumulation of an unprocessed, higher molecular mass rOGH -8 in the cytoplasmic fraction of *E. coli* cells.

### Molecular modelling analysis

In order to analyse the interaction between DsbAss peptides and Ffh M-domain, a 28 amino acid finger loop was modelled using *ab initio* method implemented in Modeller 9.15 to predict the conformation of loop region. Multiple sequence alignment was performed by HHpred between query sequence PDB ID: 1HQ1 against a database of HMMs representing proteins with known structures of Ffh M-domain of different (e.g. PBD, SCOP) or annotated protein families. It was found that Ffh M-domain of *Bacillus subtilis* (*B. subtilis*) (PDB ID: 4UE4), *Thermus aquaticus* (*T. aquaticus*) (PDB ID: 2FFH) and *Methanocaldococcus jannaschii* (*M. jannaschii*) (PDB ID: 4XCO) to be the best hits for multi-template for the disordered loop on E-value, query coverage and identity. The program employed multi-templates and modelled the missing loop of Ffh M-domain with greater than 90% confidence. Such confidence reflected the validation of core model having 2-4Å RMSD from native protein structures. Moreover, percentage identity between template and sequence greater than 40% confirmed high accuracy in model. Keeping in view the flexibility of finger loop, the model was subjected to 100ns MD simulation for structure refinement and finger loop stability. A large ensemble of 1000 snapshots generated by an MD trajectory was used to select the representative conformation from largest cluster based on RMSD cut-off of ∼1Å as shown in **Figure 4A**. This representative conformation of Ffh M-domain was further selected to perform molecular docking studies with rOGH constructs. Structural insight of the complete Ffh M-domain clearly indicated five helices (αh1-5) whose structures and spatial conformations are quite identical to those in *B. subtilis, T. aquaticus* and *M. jannaschii* [17, 23, 29]. The GTPase/M-domain linker (GM linker) together with perpendicular αh5, framed the base of groove while αh1 and the predicted finger loop (G328-D371) composed one side of the groove. All together, these structurally important segments; GM linker, αh1, finger loop, αh5 composed of hydrophobic groove which was about 12Å narrow and 16Å wide covering total area of 192Å. Within the binding groove, the hydrophobic property is maintained by D330, D333 and Q337 of αh1, M341, M344, G345, A348, S349 and M351 of finger loop, L416, F420, M423 and Q424 and M427 of αh5. 11 of these 14 residues map onto the hydrophobic countenances of α helices. The side chain flexibility of these residues, including 5 methionine, was likely to enable the hydrophobic groove to bind with signal peptides of varied lengths [30] (**Fig 4A**). The proposed signal peptide binding groove of Ffh M-domain was further examined for its size and hydrophobicity to accommodate variable signal sequences.

**Fig 4:**
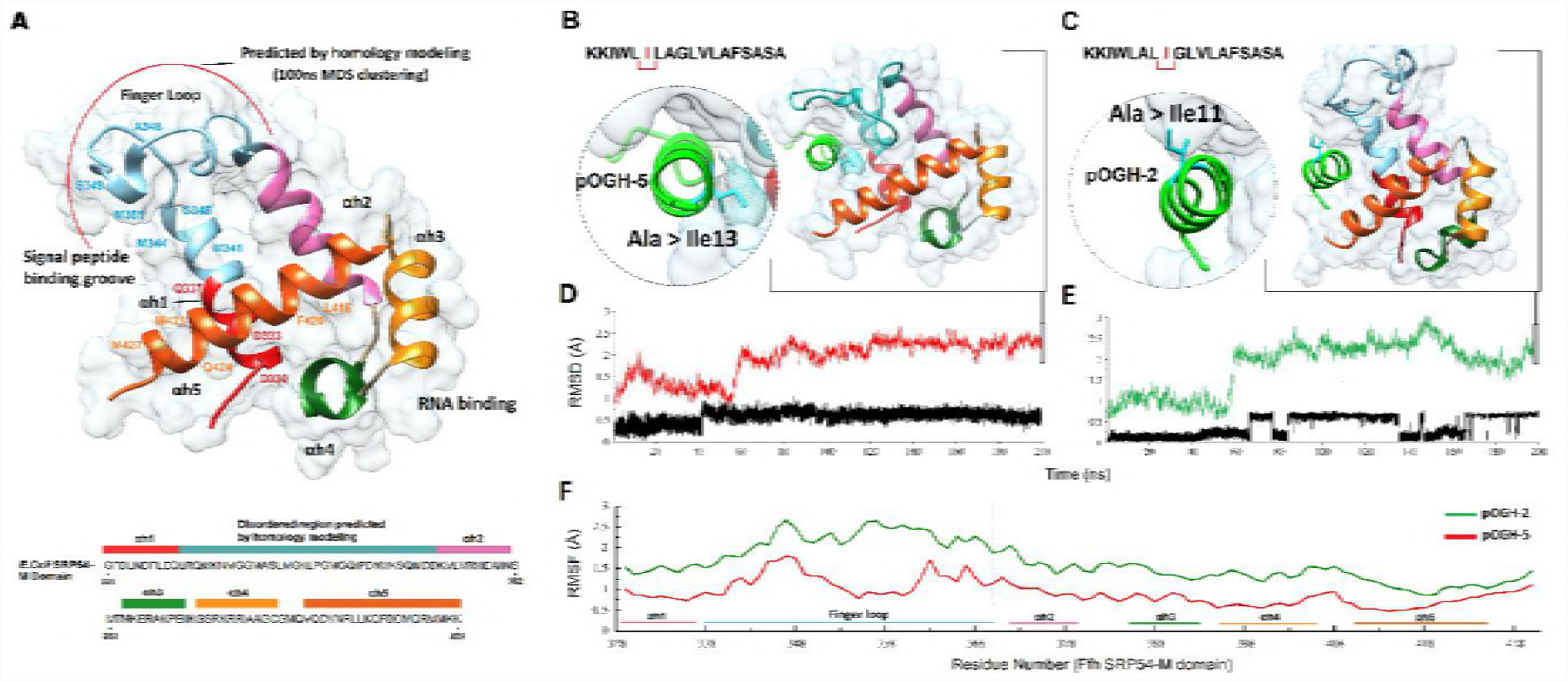
**(A)** The ribbon representation of Ffh-M-domain is enclosed by a molecular surface in white. The polypeptide chain is ramp-coloured from red (N-terminal) to orange (C-terminal). The five alpha helices are shown rendering and coloured accordingly (αh1 — red, αh2 — pink, αh3 — green, αh4 — golden, αh5 — orange). The missing disordered region near the peptide binding groove is predicted by comparative homology modelling using HHpred and coloured blue with the curved red line indicated the modelled portion. The two alpha helices (αh1 and αh5) and finger loop are making signal peptide binding groove with key residues are labelled. Right panel showed the molecular dynamic simulated complexes of M-domain with pOGH -5 **(B)** and pOGH -2 **(C)** Followed by root-mean-square-deviation trajectories in time dependent manner (ns) of pOGH -5 **(D)** pOGH -2 with red and green colour representing the M-domain and docked peptide in black. **(E)** Per-residue fluctuations of M-domain over a period of 200ns are plotted, while the corresponding helices and finger loop regions are underlined with the same colour code.

In the current study, we attempted to predict the binding mode of mutated DsbAss with Ffh M-domain. We analysed all eight DsbA constructs (pOGH -1-8) with this Ffh-M binding groove one by one and found certain alteration of amino acids in the hydrophobic part of DsbAss that underwent changes in binding with M-domain of Ffh with varying binding affinities. These alterations were carefully analysed in terms of binding energies and binding conformations of the DsbAss. For molecular docking studies, two heuristic algorithms including CLUSPRO and Patch dock were employed to examine the constancy of interaction and to locate the conformation of signal peptides in hydrophobic groove of Ffh. For the docked poses generated by Patch dock, top hundred solutions were further rescored by Fire dock algorithm, which generated the best docking model of all 8 signal peptides. To generate high quality complexes of signal peptide with Ffh, a second highly efficient protein docking algorithm CLUSPRO was employed to elucidate favourable conformations of signal peptides. The results for DsbAss 1-8, in which the lowest energy values of balanced, electrostatic, hydrophobic and VdW+Elec were highly favourable. The predicted binding energies generated by both docking algorithm are organized in **Supplementary Table S1**.

All the signal peptides underwent hydropathy analysis and results were very surprising for two constructs. The pOGH -2 and pOGH -5 constructs (both were modified with 1 Ala changed to Ile), having same hydropathy indexes (1.539) but showed varied protein expression size (∼22 and 25kDa respectively) (**Table 2**). To explore the specific role of changing Ala to IIe in directing the conformational changes, the Cα-root mean square deviation (RMSD) and root mean square fluctuation (RMSF) of Ffh M-domain complexed with pOGH -2 and -5 were calculated for the entire 200ns trajectory. RMSD clustering analysis was performed from the MD trajectory to find the most representative conformation with a cut-off value of 1.2Å to explore the binding mode of peptides with the M-domain. The conformation with a lowest RMSD was selected from the largest cluster and displayed in **Figure 4B** and **4C** and shows the RMSD of M-domain complexed with the signal peptides. With over 58ns, MD trajectories of both M-domains-peptide complexes followed a similar vertical shift of 1Å, as both have the same length of peptides. The Cα atom RMSD of M-domain-pOGH -5 showed slight deviations from 60 to 120ns followed by a stable conformation for the last 80 ns with <0.75Å deviation. Besides, the RMSD of bound pOGH -5 remained stable throughout simulation after a small shift at 42ns (**Fig 4D**). This indicated an important role in conformational transition of Ile -13 inward from the binding groove besides the finger loop adjustment and remained stable as shown in **Figure 4B**. While RMSD of Ffh M-domain showed distinct deviations after vertical shift and pOGH -2 remained unstable throughout simulation (**Fig 4D**). Additionally, the finger loop in this case, pulled to other side as the simulation progressed as compared to pOGH -5 complex (**Fig 4C**).

To analyse the mobility of each residue of M-domain around its average position with bound peptides, the RMSF values of all backbone atoms were calculated after 200ns simulations. Based on the RMSF trajectories reflected in **Figure 4F**, large fluctuations were observed in the finger loop (from Leu338 to Asp370) linking the αh1 and αh5. Among these, the largest fluctuation with pOGH -2 (**Fig 4E**) may be related to the structural shift of finger loop as compared the pOGH -5, which showed smaller fluctuation due to the stability of finger loop as shown in **Figure 4D**. The residues localized in other five helices have the smaller fluctuations with <0.5Å because of stable secondary structural elements. **Binding free energy calculations using MM-GBSA method**

### Binding free energy calculations using MM-GBSA method

To gain additional insight into the complex binding affinities, MM-GBSA analysis [31] was performed to calculate the total free energy separated by electrostatic (ΔE_ele_), van der Waals (ΔE_vdw_), solute-solvent energies (ΔG_np_ and ΔG_p_) and entropic contributions (-TΔS) as summarized in **Table 3**.

**Table 3:** MMGBSA binding free energy results for pOGH -2 and pOGH -5 (kcal/mol).

| Contributions | pOGH-2 |  | pOGH-5 |  |
| --- | --- | --- | --- | --- |
|  | Average | Std. | Average | Std. |
| $\Delta E_{\text{ele}}$ | -127.62 | 21.386 | -186.89 | 15.571 |
| $\Delta E_{\text{vdw}}$ | -91.31 | 3.708 | -100.31 | 4.341 |
| $\Delta E_{\text{MM}}$ | -218.93 | 20.924 | -287.2 | 17.986 |
| $\Delta G_{\text{p}}$ | 99.75 | 18.101 | 116.7 | 15.828 |
| $\Delta G_{\text{np}}$ | -16.66 | 0.479 | -28.05 | 0.623 |
| $\Delta G_{\text{sol}}$ | 83.09 | 18.085 | 88.65 | 15.477 |
| $\Delta G_{\text{tol}}$ | -135.84 | 4.337 | -198.55 | 4.323 |
| $-T\Delta S$ | 42.58 | 15.357 | 53.01 | 13.891 |
| $\Delta G_{\text{bind}}$ | -93.26 | | -145.54 | |

According to the **Table 3**, the overall binding free energies (ΔG_bind_) of pOGH -2 and pOGH -5 complexes are -93.26 and -145.54 kcal/mol. It should be noted that major binding free energy contributions to the peptide binding are due to the favourable electrostatic interactions for both complexes pOGH -2 and pOGH -5 (ΔE_ele_ = -127.62 and -186.86 kcal/mol) as compared to van der Waals interactions (ΔE_vdw_ = -91.31 and -100.31 kcal/mol). Whereas nonpolar solvation energies (ΔG_np_) also contributed towards peptide binding (ΔG_np_ = -16.66 and -28.05 kcal/mol) while polar solvation energy (ΔG_p_) showed unfavourable contributions. The pOGH -5 has highest ability to bind to M-domain as compared to pOGH -2 because of fairly high electrostatic binding free energy which indicated higher number of electrostatic and van der Waals interaction with M-domain.

### Per-residue decomposition analysis using MM-GBSA method

The M-domain binding groove residues that contributed to the peptide binding were further explored by per-residue decomposition analysis as illustrated in **Figure 5**. As examined, all methionine residues (M341, M344, M347, M351, Met357 and M423) of binding groove contributed strongly with both peptides. A varying trend of total interaction energies was seen in the finger loop (G328-D371) for both complexes in terms of backbone and sidechain contributions. In the case of pOGH -5, the rise in free binding energy (ΔG_bind_ = -145.54 kcal/mol) was obvious from the increased energy contribution of K353, M357, Q359, I360, D362, N363 by -8.235, -8.696, -3.328, -4.547, -2.303, -8.456 kcal/mol which were decreasing to -3.794, -0.195, -0.121, -0.809, 0.634, -0.031 kcal/mol in pOGH -2, respectively. Such increase in energy contributions was due to the structural adjustment of finger loop in a way, which positioned the sidechain conformation of some residues in favour of pOGH -5 binding as illustrated in sidechain energy percent in total interaction energy (**Fig 5**). Like M357, I360, N363, which completely oriented the sidechain position due to more stable conformation of pOGH -5-M-domain complex as shown in RMSD trajectory over a period of 200ns ( **Fig 4C** and **4D**). In contrast, these residues weakened the interaction or made them negligible as shown in **Figure 5**. Also, D422 and M423 of α5 contributed significantly with the change to Ile at position -13 in pOGH -5 with a total interaction energy of -8.569 and -9.973 kcal/mol, which is -3.882 and -5.153 kcal/mol higher than pOGH -2, whereas change of Ile to Ala at position -11 weakened the interactions due to unstable conformation of pOGH -2 (**Fig 5**). However, these important varying trends of interactions between key residues of finger loop and α5 may induce differences in the binding pose of peptides inside binding groove and overall free binding energies. The results illustrated that the change to Ile at position -11 and -13 can disrupt some interactions between the binding groove residues and peptide. Further, pOGH -5 remained stable after a small vertical shift, and structural adjustment of figure loop significantly increased the binding potential and hence could not translocate the desired rOGH into inner membrane. We found more than 60% of the protein in cytoplasmic fraction and 20% in periplasmic fraction while nothing was seen in the membrane fraction, whereas the subcellular fraction of pOGH -2 shows complete translocation of rOGH into the inner membrane (**Fig 2A**).

**Fig 5:**
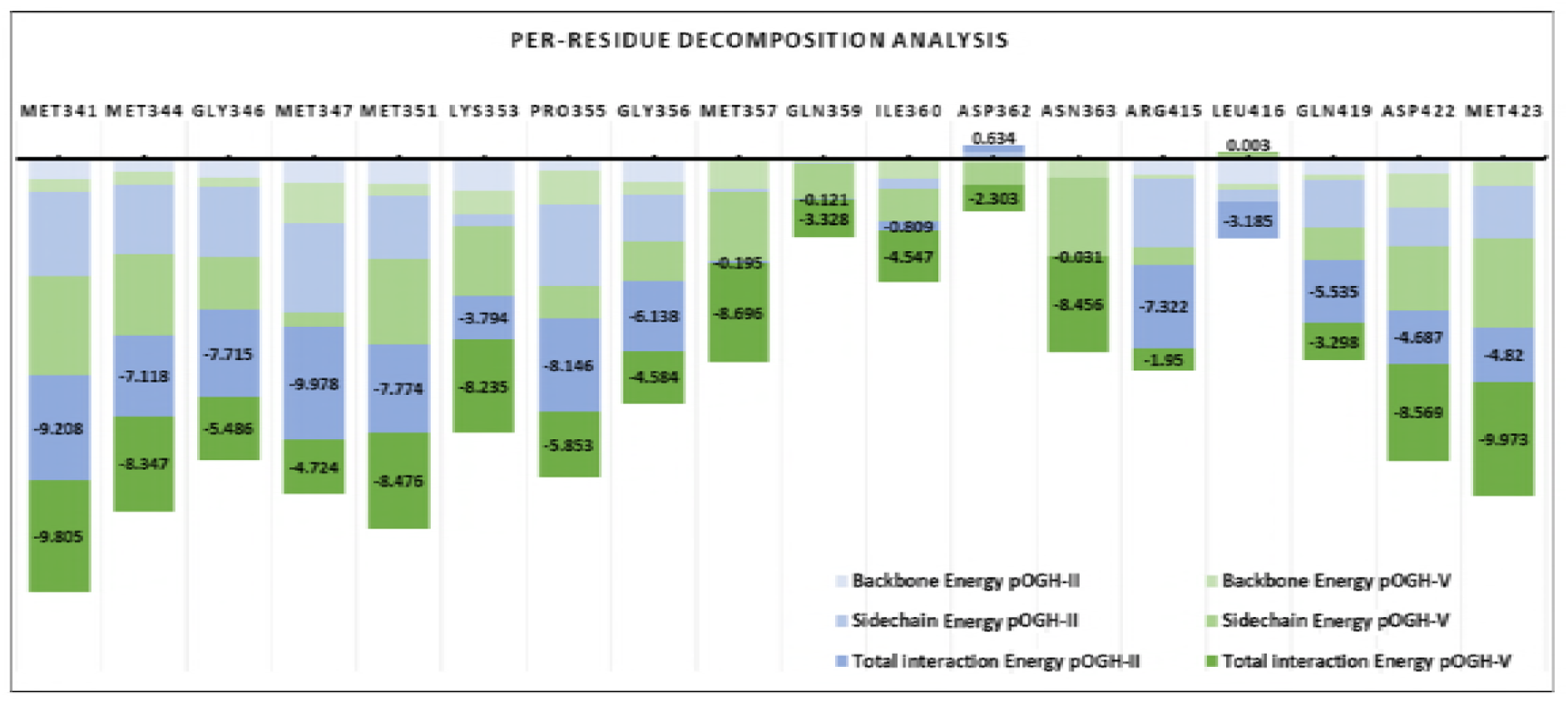
The stacking bar chart representing the binding free energy decomposition using MM-GBSA method between the pOGH -2 and pOGH -5 complexed with the M-domain. Important residues for the M-domain binding groove are labelled on the top of each bar. The total interaction energies of key residues of binding groove with pOGH -2 (blue) and pOGH -5 (green) are displayed along with sidechain and backbone contributions with same colour gradient.

### Discussion

It has been suggested by researchers that the Dsb machinery is important for the virulence factors development in the bacteria. In most cases the reduced levels of toxin subunits within cellular extracts, suggests that in the absence of DsbA catalysed disulfide bond formation the toxin subunits cannot fold into their native structures. The delivery of virulence factors can also be dependent upon DsbA-mediated disulfide bond formation. Many bacterial pathogens use T3SS to inject effector proteins into the cytosol of host cells to modulate eukaryotic cell pathways. The Dsb proteins in gram-negative bacteria aid in the maturation of exported proteins and protect them from improper oxidation [28, 32].

The proteins which are destined for extra-cytoplasmic secretion are targeted to the translocon by SRP and sec pathways. The DsbAss which we are specifically interested in, directs export through SRP mechanism. We designed a strategy to get mutation in the DsbAss and observed its effects on the translocation of rOGH as a case study. The secretion of GH in periplasmic or extra-cytoplasmic space was earlier reported [13], where DsbAss was targeted for the periplasmic secretion of human GH. Most of the work on recombinant GH is being carried out for cytoplasmic expression, for studying refolding procedures and effect of different mediums or hosts systems [33, 34]. The effect of signal peptide change, on the expression and secretion of bovine GH was studied in the past but no significant influence of the signal sequence was observed. This is the only study to our knowledge which describes the effect of specifically mutated DsbAss on the translocation of rOGH in *E. coli* cell. In the present study, we emphasized the amino acid substitution in the hydrophobic (H) part of DsbAss. It has been proven that glycine residues in the H region of GspB signal sequence affects the routing of recombinant protein towards the sec pathway [35]. In the present study, we focused mainly on the Ala residues in the hydrophobic part of DsbAss and changed each Ala in H part with the most hydrophobic amino acid Ile to observe the effect of these alteration on the translocation of rOGH. From the subcellular fraction analysis of each construct pOGH -1-8 we found interesting pattern in the two Ala positions at -11 and -13 with respect to signal peptidase site in pOGH -2 and pOGH -5 constructs respectively. The alteration of Ile in both constructs resulted in completely different translocation of rOGH; with pOGH -2, rOGH translocated to the inner membrane as reported previously for the native DsbAss [12], while pOGH -5 resulted in rOGH protein mainly stuck in the cytoplasmic fraction. This finding led us to hypothesize that there must be conformational change(s), induced by specific Ile at -13 which blocked the normal passage of translocation.

To confirm this hypothesis we performed molecular modelling analysis of these mutated protein constructs. In order to analyse the interaction between DsbAss and Ffh M-domain, a 28 amino acid residue segment was modelled and underwent structural analysis. The finger loop of Ffh has been identified to consist of conformational variability that determines its part in the binding and release mechanism of signal sequences [21, 23]. When the binding site is unoccupied, in order to make up for the hydrophobic signal sequence, the finger loop may change between conformations; open and/or closed. Thus, it potentially folds back into the groove that resides inside the Ffh crystal structure and is loaded with the finger loop of the nearby M domain [21]. All signal peptides underwent hydropathy analysis and results were very surprising for two constructs i.e. pOGH -2 and pOGH -5 which were further processed through MD simulations to analyse the conformational change with respect to backbone stability and free binding energy calculations. The most striking feature which we observed was the change in orientation of both peptides during MD simulations, where Ile of pOGH -5 is pointing in towards the groove (**Fig 4B**) and Ile of pOGH -2 is facing away from a groove (**Fig 4C**) which can eventually affect the whole mechanism of translocation. Additionally, the structural adjustment of the finger loop in a way influenced the favourable conformation of pOGH -5 inside the binding groove (**Fig 4B**). MM-GBSA calculations further elucidated the total interaction energy differences upon conformational changes. The convergence of the backbone RMSD of M-domain affirmed the stability of finger loop with pOGH -5 over time (**Fig 4D**) which anticipated high binding free energy (ΔG_bind_ = -145.54 kcal/mol) as compared to pOGH -2 (ΔG_bind_ = -93.26 kcal/mol) (**Table 3**). This high binding free energy was due to prominent electrostatic interactions which evidenced significant contributions of important finger loop residues (K353, M357, Q359, I360, D362, and N363), and were lacking in pOGH -2 (**Fig 5**). Moreover, two residues (D422 and M423) of α5 anticipated higher interaction energy -8.569 and -9.973 kcal/mol) with pOGH -5 (Ile13) and remained stable over time alongside a favourable finger loop adjustment to intact pOGH -5 inside the binding groove. The presented molecular modelling results complement with experimental findings, i.e. pOGH -5 protein construct’s subcellular fractionation showed presence of rOGH in the cytoplasmic space rather than its destination to the inner membrane as reported in our previous results [12]. The rOGH from this cytoplasmic space was biologically inactive as compared to rOGH recovered from inner membrane fraction for construct pOGH -2 was biologically active when checked on Nb2 lymphoma cells. On the basis of our findings we hypothesized a model for SRP routing of the recombinant protein through DsbAss. This means that a specific protein conformation in the hydrophobic part of the signal sequence affects binding with the M domain of the Ffh. The Ala at position -13 in the H-domain of DsbAss is important for binding of M domain of Ffh in SRP mechanism as the replacement by Ile -13 affected the interaction and changed the orientation of H part of DsbAss. As the conformation changes and regulates the latching of the signal sequence, the release of the heterodimeric domains of SRP and its receptor handover the signal sequence to the translocon [36, 37]. This change resulted in majority of the rOGH remaining in the cytoplasmic fraction without cleavage of signal peptide, as observed with the pOGH -5 construct. In order to be a biologically active rOGH, it should have disulphide bonds which is a characteristic of this protein. The mode of translocation (co-translational verses post-translational) can affect the folding process of a protein in the periplasm [28]. Therefore, we can suggest that it is not only the hydropathy but the specificity of amino acid that is important for the signal sequence in SRP mechanism. Any defect in the signal sequence can lead to improper translocation of the protein [28] which can eventually lead to formation of functionally impaired recombinant protein. We hypothesize that we can use this strategy to disarm the bacteria.

## Materials and methods

### Bacterial Strains, Plasmids and Growth Media

*E. coli* BL21 codon plus (DE3) RIPL (Stratagene, CA, U.S.A) was used for expression studies. The expression plasmid, pET22b was obtained from Novagen Inc (U.S.A) and Trizol reagent for RNA isolation from pituitary gland was purchased from Invitrogen. QIAquick gel extraction kit for DNA extraction from agarose gels and QIAprep Spin Miniprep kit for plasmid preparation were procured from QIAGEN. InsT/A cloning kit, MMLV-RTase, *Taq* DNA polymerase, restriction enzymes, IPTG and T4 DNA ligase were purchased from MBI Fermentas. Polyclonal rabbit anti-rcGH (recombinant caprine growth hormone) antibody was used from the in house production facility at School of Biological Sciences, University of the Punjab, Lahore, Pakistan.

### Total RNA Isolation and cloning of OGH gene

Total RNA was isolated from pituitary gland of local ovine breed, “LOHI” using Trizol reagent. RT-PCR was performed using total RNA as template and the primers were designed on the basis of OGH sequence. The forward primer (FP); FP-1 (5`ATC CAT GGC CTT CCC AGC CAT G3`) and the reverse primer (RP); RP-1 (5`TAG GAT CCG CAA CTA GAA GGC AGC 3`) contained *Nde*I and *Bam*HI restriction sites at the 5’end terminal respectively. M-MuLV RTase was used to catalyse the reverse transcription reaction and the synthesized cDNA was amplified using the above set of primers. The amplified product was ligated into the cloning vector pTz57R/T by T/A cloning technique to produce the recombinant pTzOGH-1 construct. PCR amplification of the first strand reaction by *Taq* DNA polymerase was carried out in Fermentas Biosystems 2720 thermal cycler by denaturation, annealing and extensions at 94°, 65° and 72°C, respectively with a hold time of 1 minute each for 28 cycles.

### Construction of Recombinant Expression Plasmids

Eight constructs based on the DsbAss variation upstream of the 5’-start codon of OGH gene were produced. These were studied in order to observe the expression and proper translocation of OGH into the extra-cytoplasmic space of *E. coli.* The eight FP; FP-1 to FP-8 were designed with *Nde*I site at 5’ end, 57 nucleotides of the DsbAss (underlined) and six nucleotides of the OGH gene (bold) as shown in **Table 1**. The FP-1 had native DsbAss, however the rest of the FP-2 to FP-8 were designed with the change of amino acid in H, C and N domains of DsbAss (**Table 1)**. Only one reverse primer was used i.e. RP1 (5`TAG GAT CCG CAA CTA GAA GGC AGC 3`) with *Bam*HI site at the 5’-end. The wild-type GH sequence in the construct pTzOGH-1 was amplified using each of the forward (FP-1 to FP-8) and the reverse primer. These PCR products and pET22b were then digested with *Nde*I/*Bam*HI and ligated thus to generate a series of recombinant plasmids (pOGH -1-8). These constructs were primary cloned in *E. coli* strain, DH5 and further into BL21 codon plus for the expression studies.

### Expression and sub-cellular fractionation of pOGH constructs

The cells from a single colony of each of the transformants of all the constructs were transferred to LB medium (100μg/ml ampicillin) and grown overnight at 37°C in an orbital incubator shaker at 150rpm. Next morning 100ml LB medium was inoculated with 1% (v/v) of overnight culture for each construct and induced with 10μM IPTG at 0.5 OD_600_. After 6hrs post induction at 37°C, 150rpm shake rate, cells were harvested by centrifugation at 5,000 x *g* for 5 minutes. The cell pellet was resuspended in 20ml of STE buffer [(20% v/v) sucrose in 0.33 M Tris-HCl, 1mM EDTA (pH8.0)] and left at room temperature for 10 minutes and then centrifuged at 7,500 x *g* for 10 minutes at 4°C. Most of the supernatant was discarded while leaving behind approximately 250μl volume to initially resuspend the cells and the proteins in the periplasm were extracted by rapid addition of 10ml of chilled 0.5mM MgCl2 using osmotic shock [38]. The cell suspension was left on ice for 10 minutes and the periplasmic fraction was collected after centrifugation at 5,000 x *g* for 5 minutes at 4°C. The cell pellet was processed further for the cytoplasmic and membrane fractions by re-suspension in 20ml TE buffer [0.3M Tris-HCl, 1mM EDTA (pH 8.0)] and sonication by 6 bursts of 30 seconds each with 3 minute interval. The sonicated sample was centrifuged at 5,000 x *g* for 20 minutes at 4°C and the supernatant was retained as the cytoplasmic fraction. The pellet was washed with 5ml 0.3M sucrose, 0.3M Tris-HCl (pH8.0) and centrifuged at 11,000 x *g* for 10 minutes. The supernatant was ultra-centrifuged at 100,000 x *g* for 1hr and the membrane pellet was finally resuspended in 5ml of 0.3M Tris-HCl (pH8.0) [38].

### Protein and Western blot analysis

Protein analysis of the lysates were carried out on 15% SDS-PAGE according to the method described by Laemmli [39]. Resolved proteins were visualized by staining with Coomassie brilliant blue R250. The relative amount of protein in the bands were detected by using BioRad GS-800 calibrated Densitometer and quantified by Phoretix 1Dsoftware (version 5.1) operating under MS Windows XPTM. Protein content was estimated spectrophotometrically at 595 nm with Coomassie Blue G-250 dye-binding procedure of Bradford assay [40], employing bovine serum albumin (BSA) as the standard.

Further confirmation for the identity of the membrane bound rOGH protein was carried out by Western blot analysis using polyclonal rabbit anti-rcGH antibody derived from recombinant caprine GH which confirmed it as GH.

### Molecular modelling studies

The Ffh M domain was retrieved from Protein Data bank under PDB ID: 1HQ1 (Resolution: 1.52, R-Free: 0.199) [41]. The co-crystallized structure with RNA was carefully investigated and revealed the finger loop, a 28-amino acid residue segment of M domain containing the proposed signal peptide recognition site is disordered in the crystals and was not observed in the electron density map. Therefore homology modelling was performed for this segment by taking Ffh SRP54 domains from different organisms *B. subtilis* (PDB ID: 4UE4), *T. aquaticus* (PDB ID: 2FFH) and *M. jannaschii* (PDB ID: 4XCO). HHpred (Homology detection & structure prediction by HMM-HMM comparison) was employed to search for possible templates and multi-template homology modelling was performed by using MODELLER v9.15 [42, 43]. The stereochemical assessment of modelled loop and residue-by-residue geometry were checked by PROCHECK [44] and Molprobity [45]. Due to the flexibility of finger loop, the modelled M domain was executed through 100 ns MD simulations using AMBER 16 to optimize a most stable conformation using RMSD clustering (Supplementary information for detailed MD protocol). In order to analyse the DsbAss conformation inside the signal peptide groove, protein molecular docking protocols were employed to obtain the most favourable conformation followed by MD simulations. PatchDock [46] with rescoring algorithm FireDock server [47] and Cluspro v2 [48] were used for molecular docking. Input parameters for PatchDock tool were PDB 3D coordinate files with default parameters. To evaluate each candidate transformation, scoring function was used that considered to have both atomic desolvation energy and geometric fit. Finally, clustering based on Root mean square deviation (RMSD) was applied to each solution to discard the redundant solution.

ClusPro v 2.0 is the first fully automated and top ranked performance in the latest rounds of CAPRI experiments [49]. It evaluates the putative structure based on good electrostatic and desolvation free energies for clustering and rank them accordingly. ClusPro server produce four categories of predictive models which are ranked by cluster size including, balanced, electrostatic favoured, Hydrophobic favoured and VdW + Elec. For the current work, saturated clusters of best models were focused in all categories. The binding energies were calculated and protein complexes of Ffh M-domain with all DsbAss were visually inspected for binding site residues and molecular interactions using UCSF Chimera v10 [50]. Binding site residues were defined, as those residues with at least one heavy atom inside 4Å distance from any heavy atom of the binding groove. To check the role in changing Ile to Ala at position -11 and -13 in pOGH -2 and pOGH -5 respectively, an extensive MD simulations of 200 ns were carried out to better comprehend the conformational stability of the DsbAss in signal peptide binding groove. All MD simulations were executed using the AMBER 16 package [51]. After a stepwise minimization, heating and equilibration protocol, the solvated system with TIP3P water molecules was subjected to a production simulation of 200 ns at 300K and at 1 bar pressure. The detailed MD simulation protocol is described in supplementary information (**Supplementary Information**). The cpptraj module of AMBER 16 [52] was used for the trajectory analysis.

### MM-GBSA binding free energy calculations

The binding free energy (ΔG_bind_) of pOGH -2 and pOGH -5 complexes were calculated through MM-GBSA module of AMBER 16. The details of the MM-GBSA method have been explained elsewhere [53, 54]. The snapshots from last 20 ns MD trajectory of the complex with an interval of 2.0 ps were generated and binding free energy (ΔG_bind_), which is the sum of molecular mechanics energy (ΔEMM), solvation free energy (ΔG_sol_) and entropic (-TΔS) contribution. Whereas both ΔGMM and ΔGsol are further divided into internal energy (ΔE_int_), electrostatic (ΔE_ele_), van der Waals (ΔE_vdw_) energy in the gas phase, and polar (ΔGp) and non-polar (ΔG_np_) contributions to the solvation free energy as follows:

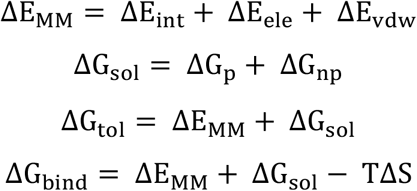

Normal mode analysis (nmode module) of AMBER 16 [51] was performed with MM-GBSA method to calculate the contributions of each residue of binding groove to the total binding free energy.

## ACKNOWLEDGEMENTS

This study was supported by a grant from Higher Education Commission, Government of Pakistan. The authors greatly acknowledge the valuable inputs and guidance of late Dr. Mushtaq Kaderbhai in this study.

